# Experimental infection of cattle with *Mycobacterium tuberculosis* isolates shows the attenuation of the human tubercle bacillus for cattle

**DOI:** 10.1101/202663

**Authors:** Bernardo Villarreal-Ramos, Stefan Berg, Adam Whelan, Sebastien Holbert, Florence Carreras, Francisco J. Salguero, Bhagwati L. Khatri, Kerri M. Malone, Kevin Rue-Albrecht, Ronan Shaughnessy, Alicia Smyth, Gobena Ameni, Abraham Aseffa, Pierre Sarradin, Nathalie Winter, Martin Vordermeier, Stephen V. Gordon

## Abstract

The *Mycobacterium tuberculosis* complex (MTBC) is the collective term given to the group of bacteria that cause tuberculosis (TB) in mammals. It has been reported that *M. tuberculosis* H37Rv, a standard reference MTBC strain, is attenuated in cattle compared to *Mycobacterium bovis*. However, as *M. tuberculosis* H37Rv was isolated in the early 1930s, and genetic variants are known to exist, we sought to revisit this question of attenuation of *M. tuberculosis* for cattle by performing a bovine experimental infection with a recent *M. tuberculosis* isolate. Here we report infection of cattle using *M. bovis* AF2122/97, *M. tuberculosis* H37Rv, and *M. tuberculosis* BTB1558, the latter isolated in 2008 during a TB surveillance project in Ethiopian cattle. We show that both *M. tuberculosis* strains caused reduced gross and histopathology in cattle compared to *M. bovis*. Using *M. tuberculosis* H37Rv and *M. bovis* AF2122/97 as the extremes in terms of infection outcome, we used RNA-Seq analysis to explore differences in the peripheral response to infection as a route to identify biomarkers of progressive disease in contrast to a more quiescent, latent infection. Our work shows the attenuation of *M. tuberculosis* strains for cattle, and emphasizes the potential of the bovine model as a ‘One Health’ approach to inform human TB biomarker development and post-exposure vaccine development.

## Introduction

The *Mycobacterium tuberculosis* complex (MTBC), the group of pathogens that cause tuberculosis (TB) in mammals, show distinct host preference. *Mycobacterium tuberculosis* is the hallmark member of the MTBC and the most deadly human pathogen globally, with close to 2 billion people infected worldwide ^1^. The animal-adapted members of the MTBC are named after the host of initial/most frequent isolation, and comprise: the exemplar animal pathogen and predominant agent of bovine TB *M. bovis* ^2^; *M. microti* ^3^; the ‘Dassie bacillus’ ^4,5^; *M. caprae* ^6^; *M. pinnipedii* ^7^; *M. orygis* ^8^, and *M. mungi* ^9^. A caveat is that these species designations do not define host exclusivity; MTBC members can infect a range of mammals to greater or lesser degrees. The central feature of host adaptation is the ability to sustain within a host population. Thus, *M. bovis* can infect and cause disease in humans; however, the capacity of *M. bovis* to transmit between immunologically competent humans is severely limited compared to *M. tuberculosis* ^10,11^. Similarly, reports suggest that *M. tuberculosis* appears attenuated in a bovine host compared to *M. bovis* ^12–14^.

The advent of systems for mutagenesis of mycobacteria allied with (largely) murine screens for phenotype has provided enormous insight into the virulence genes and pathogenic strategies of mycobacteria. Yet the basis for host preference across the MTBC is largely unknown. Members of the MTBC share >99% nucleotide identity across their genomes, with for example ~2,400 SNPs between *M. bovis* AF2122/97 and *M. tuberculosis* H37Rv ^15–17^. Regions of difference (RD), deleted loci absent from one MTBC member relative to another, serve as unique markers to differentiate species with some having had functional roles ascribed. The RD1 locus of *M. tuberculosis* encodes a type VII secretion system whose role in virulence of *M. tuberculosis* has been convincingly explored using a range of *in vitro* and model systems ^18^. However, RD1-like deletions from *M. microti* and the ‘dassie’ bacillus indicate that discrete evolutionary scenarios have moulded the virulence strategies and genomes of the MTBC bacilli ^19,20^. Similarly, functional impacts of single nucleotide polymorphisms (SNPs) between the various MTBC and their potential role in host-pathogen interaction have been described ^21^. Much work remains in describing the precise host and pathogen molecular factors involved in host preference ^22^.

An essential step to defining host tropic factors is defining tractable experimental models; given the range of wild and domesticated mammals that are susceptible to infection by MTBC members, this is no small task. Nevertheless, with the aim of exploring MTBC host preference, we have previously explored the comparative virulence of *M. tuberculosis* and *M. bovis* in a bovine experimental infection model, and showed that *M. tuberculosis* H37Rv was attenuated compared to *M. bovis* AF2122/97 ^13^. However, while *M. tuberculosis* H37Rv is the reference strain of the MTBC, its isolation was first reported 1935 ^23^ and it has been maintained in multiple laboratories globally, with separate extant ‘versions’ of *M. tuberculosis* H37Rv ^24^. The possibility remained that other *M. tuberculosis* clinical isolates would show a different phenotype in a bovine infection. Indeed, there have been increasing numbers of reports of the isolation of *M. tuberculosis* from cattle ^25–27^, including our own work where we previously isolated *M. tuberculosis* strains from lesions identified in cattle at slaughter in Ethiopian cattle ^28,29^. This would suggest that either there exist strains of *M. tuberculosis* with virulence characteristics that allow them to infect and cause disease in cattle, or that the cattle from which these *M. tuberculosis* strains were isolated had greater susceptibility to infection due to being immune comprised, co-infections, age, malnutrition, or other predisposing factors such as being in an environment of continuous exposure to *M. tuberculosis*.

We therefore set out to evaluate the attenuated virulence of *M. tuberculosis* in the bovine host using a recent *M. tuberculosis* bovine isolate as the comparator. We chose to use an Ethiopian *M. tuberculosis* strain that had been isolated from a Zebu bull, *M. tuberculosis* BTB1558, to perform this new experimental infection in cattle, and to compare the virulence of this latter isolate to the *M. tuberculosis* H37Rv and *M. bovis* AF2122/97 strains.

## Material and Methods

### Ethical permission

Ethical permission was obtained from the APHA Animal Use Ethics Committee (UK Home Office PCD number 70/6905), AHRI/ALERT Ethics Review Committee (Ethiopia) and the French Research and Education Ministry, via the Val de Loire Ethical Committee (CEEAVDL, #19) for INRA (France).

### Mycobacterial strains and culturing protocols

Three strains were used for this bovine challenge experiment: *M. bovis* AF2122/97 is a field strain isolated from a cow in Great Britain in 1997 ^16^. *M. tuberculosis* H37Rv was from the APHA culture stocks. The virulence of both the *M. bovis* AF2122/97 and *M. tuberculosis* H37Rv stocks has been confirmed via inoculation of guinea pigs ^13^, with both seed stocks clearly virulent in this model. *M. tuberculosis* BTB1558 was isolated in 2008 from the cranial mediastinal lymph node of a Zebu bull (*Bos indicus*) at Ghimbi abattoir, Ethiopia. The lesion from which the strain was isolated was classed as localised, and was not caseous or calcified; no nasal secretions were present at the ante-mortem investigation. An *M. tuberculosis* strain of the same spoligotype as BTB1558 (SIT 764) was isolated from a human pulmonary TB patient in Ethiopia ^30^; both the cattle and the human isolate have been confirmed as members of the Euro-American lineage, also known as Lineage 4 ^31,32^.

*M. bovis* AF2122/97 and *M. tuberculosis* BTB1558 had been passaged a maximum of five times prior to the challenge experiment. Seed stocks had been cultured to mid-log phase in Middlebrook 7H9 media (Difco, UK) supplemented with 10% (v/v) Middlebrook acid-albumin-dextrose-catalase enrichment (Difco), 4.16 g/l sodium pyruvate (Sigma-Aldrich, UK) and 0.05% (v/v) Tween 80 (Sigma-Aldrich) and stocks stored frozen at −80°C. The colony forming units (CFU)/ml of bacterial stocks, infection inoculum, and homogenised tissue was determined by bacterial enumeration of a serial dilution cultured on modified Middlebrook 7H11 agar ^33^. Inoculated plates were incubated at 37°C for four weeks (six weeks for tissue samples) prior to enumeration of individual colonies on the agar plates. All seed stocks were confirmed with a viability of approximately 2 × 10^7^ CFU/ml prior to further use.

### Preparation of M. bovis and M. tuberculosis infection inoculum

Frozen seed stocks were thawed and diluted in 7H9 medium to a final concentration of approximately 5 × 10^3^ CFU/ml. For each animal, 2ml of this infection inoculum were drawn into a 5ml Luer-lock syringe. An aliquot of the *M. bovis* AF2122/97, *M. tuberculosis* H37Rv, and *M. tuberculosis* BTB1558 inocula was retained to determine, retrospectively, the concentration of bacilli used in each inoculum. Heat-inactivated samples of each strain were used to identify the strains by large sequence polymorphism ^29,34^ and spoligotyping ^35^.

### Cattle infection

Twelve female Limousin x Simmenthal cattle of approximately six months of age raised from birth within INRA’s animal facility were divided into three groups of four. An infective dose of 1×10^4^ CFU was targeted for each strain; inocula were delivered endobronchially in 2 ml of 7H9 medium as described previously ^36^. In brief, animals were sedated with xylazine (Rompun^®^ 2%, Bayer, France) according to the manufacturer’s instructions (0.2 mL/100 kg, IV route) prior to the insertion of an endoscope through the nasal cavity into the trachea for delivery of the inoculum through a 1.8 mm internal diameter cannula (Veterinary Endoscopy Services, U.K.) above the bronchial opening to the cardiac lobe and the main bifurcation between left and right lobes. Two ml of PBS were used to rinse any remains of the inoculum into the trachea and then cannula and endoscope were withdrawn. The canal through which the cannula was inserted into the endoscope was rinsed with 20 ml of PBS and the outside of the endoscope was wiped with sterilizing wipes (Medichem International, U.K.) prior to infection of the next animal. Retrospective counting of the inocula revealed infection with 1.66×10^4^ CFU *M. tuberculosis* H37Rv; 2.2×10^4^ CFU *M. tuberculosis* BTB1558; and 1.12×10^4^ CFU *M. bovis* AF2122/97.

### Monitoring of infection by the IFN-γ release Assay (IGRA)

Blood was collected from animals one day prior to the infectious challenge (-1) and every two weeks after infection until the animals were culled. Heparinized whole blood (250μl) was incubated with a selection of mycobacterial antigens: PPD-Avium (PPD-A) or PPD-Bovine (PPD-B) (Prionics) respectively at 25 IU and 30 IU final; or peptide pools covering ESAT6/CFP10, Rv3873 or Rv3879c added in a volume of 25μl to a final concentration of 10μg/ml. Pokeweed mitogen was used as the positive control at 10μg/ml, and a media-only negative control also included. After 24h in 5% (v/v) CO_2_ atmosphere at 37°C stimulated bloods were centrifuged (400xg for 5 min); 120μl of supernatant was removed and stored at −80°C for subsequent IFN-γ quantification using the Bovigam kit (Prionics) in accordance with the manufacturer’s instructions.

For RNA extractions, 4 ml of heparinized whole blood was incubated with PBS or PPD-B in the same condition as described above. After 24 hr 3ml of stimulated blood were transferred to Tempus™ Blood RNA tubes (Life Technologies) and stored at −80°C. The remaining 1ml was centrifuged and the supernatant stored at −80°C for cytokine analyses.

### Multiplex ELISA analyses

PPD-B stimulated whole blood supernatants were assayed for cytokine levels using a custom-designed bovine Meso Scale Discovery^®^ (MSD) multiplex protein analysis platform (Meso Scale Discovery^®^, Gaithersburg, MD, USA). The bovine cytokines analysed were: IL-1β, IL-6, IL-10, IL-12, and TNF-α. Multiplex 96 well plates (supplied with target capture antibodies spotted onto separate carbon electrodes in each well) were blocked with MSD^®^ assay buffer for 30 min at room temperature before the addition of 25μl samples or MSD^®^ standards (prepared according to manufacturer’s instructions). Following 2h sample incubation at room temperature, plates were washed and incubated for a further 2h with a combined cocktail of biotinylated detection antibodies for each cytokine and MSD^®^ SULFO-TAG™-labelled Streptavidin (according to the manufacturers’ instructions). After a final wash, plates were coated with MSD^®^ Buffer-T and luminescence (OD_455nm_) was measured on a SECTOR^®^ Imager 6000 instrument (MSD). IL-6, IL-10 and IL-12 responses are reported as U/ml while IL-1β and TNF-α responses are reported in ng/ml as interpolated from the standard curves for each cytokine included on each plate.

### Gross pathology and histopathology

Ten weeks after infection animals inoculated with *M. tuberculosis* were killed and subjected to post-mortem analysis as indicated elsewhere ^37^; animals inoculated with *M. bovis* were sacrificed six weeks after infection for post-mortem analysis as above. The presence of gross pathological TB-like lesions was scored as previously described (37). For histology, a cross-sectional slice of the lymph node was collected into a 100 ml pot containing buffered formalin. Collected tissue samples were shipped overnight from INRA to APHA Weybridge for subsequent processing.

Tissues evaluated for gross pathology included the following lymph nodes: left and right parotid, lateral retropharyngeal, medial retropharyngeal, submandibular, caudal, cranial mediastinal and cranial tracheobronchial and pulmonary lymph nodes; lung tissue samples were also taken. Tissue samples were preserved in 10% phosphate buffered formalin for 7 days before being embedded in paraffin wax. Four-micron sections were cut and stained with hematoxylin and eosin (H&E); Ziehl-Neelsen staining was carried out for the detection of acid-fast bacilli (AFB). For histopathology, sections of thoracic (caudal mediastinal, cranial mediastinal, cranial tracheobronchial, left and right bronchial) and extrathoracic (left and right parotid, left and right medial retropharyngeal, left and right lateral retropharyngeal, left and right mandibular) lymph nodes, left and right tonsils and lung were stained with for examination by light microscopy to assess the number, developmental stage and distribution of each granuloma (I-IV) as well as assessing the quantity and location of AFB as previously described ^38,39^.

### Evaluation of tissue bacterial load

For bacteriology, up to 20 g of tissue was collected into 25 ml sterile containers and frozen at −80°C on the day of collection. Frozen tissues were shipped at +4°C to APHA Weybridge and immediately upon arrival were homogenised using a Seward Stomacher Paddle Blender with bacterial enumeration undertaken as previously described ^37^. Macerates were plated on modified 7H11 agar plates containing 10% (vol/vol) Middlebrook oleic acid-albumin-dextrose-catalase enrichment ^33^. Plates were seeded with 500μl, 50μl and 5μl of macerate; 450μl and 500μl of PBS was added to the plates containing 50μl and 5μl respectively to help distribute the macerate on the whole plate. Using this method the limit of detection was 2 CFU/ml.

### RNA extraction and library preparation

Thirty-two strand-specific RNA-Seq libraries were prepared from whole blood from *M. bovis* AF2122/7 and *M. tuberculosis* H37Rv infected animals (n=4) at day 14 and day 42 that were either stimulated or not with PPD-B. Total RNA including miRNA was extracted from the Tempus™ Blood RNA Tubes using the Preserved Blood RNA Purification Kit I (Norgen Biotek Corp, Canada) according to the manufacturer’s instructions. Twelve random samples were chosen to assess RNA integrity using the RNA 6000 Nano Kit (Agilent) in conjunction with the Agilent 2100 Bioanalyzer. RNA Integrity Numbers (RINs) ranged from 8 to 9.1 (8.6 average). RNA was quantified using the NanoDrop™ ND-1000 Spectrophotometer (Thermo Fisher Scientific) and 200ng of total RNA was subjected to two rounds of Poly(A) mRNA purification using the Dynabeads® mRNA DIRECT™ Micro Kit (Invitrogen™) according to the manufacturer’s recommendations. The samples were prepared for sequencing using the ScriptSeq™ v2 RNA-Seq Library Preparation Kit, Index PCR Primers and the FailSafe™ PCR enzyme system according to manufacturer’s specifications (Illumina® Inc., Madison, WI, USA). The Agencourt® AMPure® XP system (Beckman Coulter Genomics, Danvers, MA, USA) was utilised to purify the resulting RNA-Seq libraries. Libraries were quantified using the Quant-iT dsDNA Assay Kit and subsequently pooled in equimolar concentrations (Thermo Fisher Scientific). The 32 sequencing libraries were pooled and sequenced over three lanes of an Illumina HiSeq 2500 Rapid Run flow cell (v1) in paired end (2 x 100 bp) format by Michigan State University Research Technology Support Facility, Michigan, USA. Base-calling and demultiplexing was performed with Illumina Real Time Analysis [v1.17.21.3] and Illumina Bcl2Fasta [v1.8.4] respectively. The RNA-Seq data has been deposited in the European Nucleotide Archive, accession number PRJEB22247.

### RNA-Seq pipeline

The pipelines used for the analysis of the RNA-Seq data are available on GitHub (https://github.com/kerrimalone). Computational analyses were performed on a 32-node Compute Server with Linux Ubuntu [version 12.04.2]. Briefly, pooled libraries were deconvoluted, adapter sequence contamination and paired-end reads of poor quality were removed using cutadapt [v1.13] (Phred > 28) ^40^ and the filterbytile.sh script from the BBMap package ^41^. At each step, read quality was assessed with FastQC [v0.11.5] ^42^. Paired-end reads were aligned to the *Bos taurus* reference genome *(B. taurus* UMD 3.1.1) with the STAR software ^43^. Read counts for each gene were calculated using featureCounts, set to unambiguously assign uniquely aligned paired-end reads in a stranded manner to gene exon annotation *(B. taurus* UMD 3.1.1 GCF_000003055.6) ^44^ Differential gene expression analysis was performed using the edgeR Bioconductor package that was customised to filter out all bovine ribosomal RNA (rRNA) genes, genes displaying expression levels below one count per million (CPM) in at least four individual libraries and identify differentially expressed (DE) genes correcting for multiple testing using the Benjamini-Hochberg method with a log2 fold change (log2FC) greater than 1/less than −1 and a False-Discovery Rate (FDR) threshold of significance ≤ 0.05 ^45^. DE gene expression was evaluated between *M. bovis* and *M. tuberculosis* infected animals for unstimulated blood samples (unpaired statistics) for each time point in addition to between unstimulated and PPD-B-stimulated blood samples for the *M. bovis* and *M. tuberculosis* infected animals independently at each time point (paired statistics). Cellular functions and pathways over-represented in DE gene lists were assessed using the SIGORA R package while graphical representation of data results was achieved using the R packages ggplot2, VennDiagram and related supporting packages ^46–48^.

### miRNA RT-qPCR

MicroRNA miR-155 was selected for analysis based on its suggested role in immune response to *M. bovis* infection ^49^. As the human and bovine sequences are identical, hsa-miR-155-5p primers were purchased from Exiqon miRCURY UniRT miRNA primers (catalogue number 204308).

## RESULTS

### Immune response in infected cattle

Three groups of four cattle each were infected endobronchially with 1.66×10^4^ CFU *M. tuberculosis* H37Rv, 2.2×10^4^ CFU *M. tuberculosis* BTB1558, or 1.12×10^4^ CFU *M. bovis* AF21222/97, respectively. Because of the need to restrict the total time the experiment would run in the containment facility, animals infected with *M. tuberculosis* strains H37Rv or BTB1558 were maintained for 10 weeks, while *M. bovis* AF2122/97-infected animals were maintained for 6 weeks, after which all animals underwent post-mortem examination. Antigen specific IFN-γ responses were detected against both PPD-B (data not shown) and a cocktail of ESAT-6/CFP-10 peptides (Figure 1) two to three weeks after infection, with no significant difference in responses between groups over the course of the infections.

**Figure 1:**
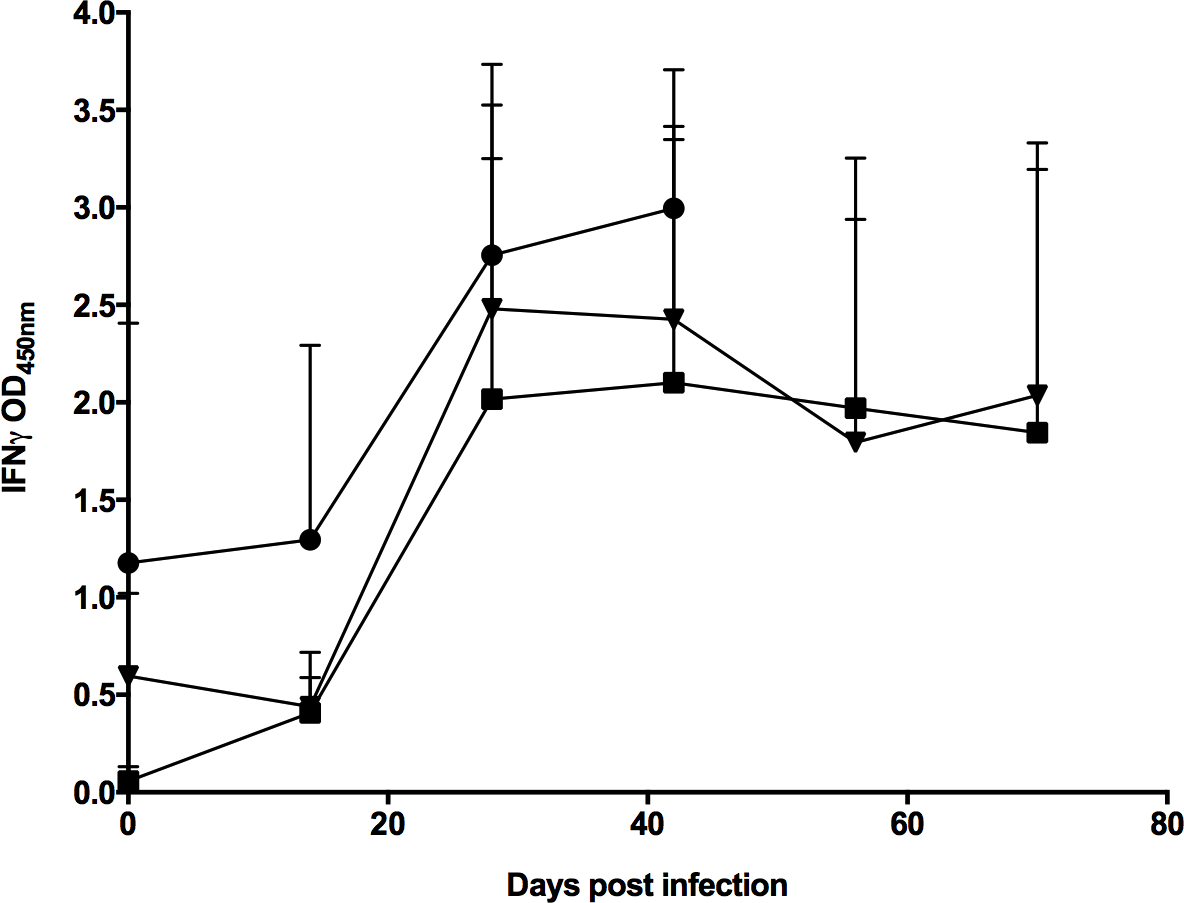
Infection of cattle with *M. tuberculosis* H37Rv, *M. tuberculosis* BTB1558 or *M. bovis* AF2122/97 induces similar peripheral immune responses. Blood was collected at regular intervals from cattle prior to and after experimental infection with *M. tuberculosis* H37Rv (n=4), *M. tuberculosis* BTB1558 (n=4), or *M. bovis* AF2122/97 (n=4). Whole blood was isolated and stimulated with a cocktail of peptides derived from ESAT-6 and CFP-10. The responses in the infected cattle are shown: *M. tuberculosis* H37Rv (squares); *M. tuberculosis* BTB1558 (triangles); *M. bovis* AF2122/97 (circles). *M. bovis* infected animals were maintained for 6 weeks, *M. tuberculosis* groups for 10 weeks. Data for each time point is presented as the mean response ± SEM.

In previous work ^13^ we had seen that while stimulation of whole blood with the antigen Rv3879c triggered IFN-γ production in *M*. bovis-infected cattle, *M. tuberculosis* H37Rv-infected cattle showed no responses. In this current work Rv3879c stimulation of whole blood provided less definitive outcomes, as while Rv3879c triggered minimal IFN-γ responses in blood from *M. tuberculosis* H37Rv or BTB1558 infected animals, the baseline responses to Rv3879c stimulation in *M. bovis* AF2122/97 infected cattle were high prior to infection, and showed no increase in response over the course of infection (data not shown).

Supernatants from PPD-B stimulated samples were also checked for IL-1β, IL-6, IL-10, IL-12, and TNF-α production over the course of infection using the MSD multiplex platform (Figure 2). Over the first 6 weeks after infection, all animals showed similar responses to all strains, although *M. tuberculosis* BTB1558 generated higher IL-1β responses at day 28 compared to responses induced by *M. tuberculosis* H37Rv or *M. bovis* AF2122/97. *M. tuberculosis* BTB1558 induced higher production of IL-10, IL-12, and TNF-α than *M. tuberculosis* H37Rv at the later time points (days 56 and 70) due to responses waning in the *M. tuberculosis* H37Rv group from day 42 onwards. Less than 1U/mL of IL-6 were detected in both infection groups (data not shown).

**Figure 2:**
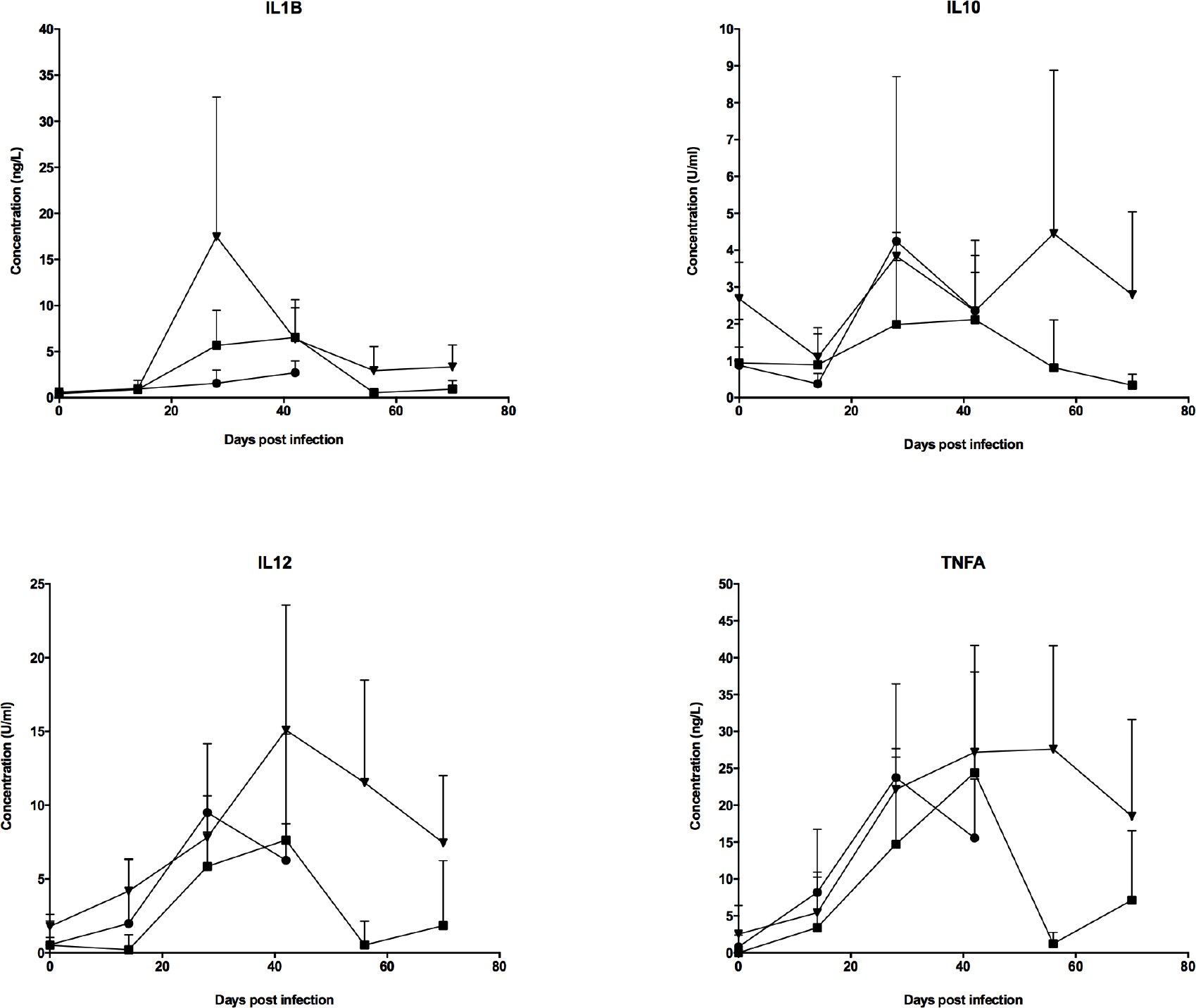
Cytokine analysis of stimulated whole blood from *M. tuberculosis* H37Rv, *M. tuberculosis* BTB1558 or *M. bovis* AF2122/97 infected cattle. PPD-B stimulated-whole blood supernatants were assayed for IL-1β, IL-10, IL-12 (D), and TNF-α cytokine levels using a custom-designed bovine MSD. The response in the *M. tuberculosis* H37Rv infected cattle is shown as squares; *M. tuberculosis* BTB1558 is shown as triangles; and *M. bovis* AF2122/97 shown as circles; all groups contained 4 animals. IL-10 and IL-12 responses are reported as U/ml while IL-1β and TNF-α responses are reported in ng/ml as interpolated from the standard curves for each cytokine included on each plate. Data for each time point is presented as the mean response ± SEM.

### Gross pathology

As described above, animals infected with *M. bovis* AF2122/97 were culled 6 weeks post-infection, while *M. tuberculosis* groups were culled after 10 weeks. Lungs and lymph nodes (thoracic and extrathoracic) were examined for gross lesions. Typical gross lesions were isolated granulomas or coalescing clusters of granulomas of variable size ranging from 5 to 10 mm in diameter. Gross pathology scores are summarised in Figure 3(A-D). Statistically significant differences were observed in lung gross lesions of animals infected with *M. bovis* compared to animals infected with *M. tuberculosis* H37Rv or *M. tuberculosis* BTB1558; no significant differences were observed comparing the lung lesions of the two sets of *M. tuberculosis* infected animals. Lung gross lesions were only observed in animals infected with *M. bovis* AF2122/97 with scores ranging from 5 to 10 and scores of 0 (no visible lesions) in animals infected with *M. tuberculosis* H37Rv or BTB1558. Thoracic lymph nodes (cranial mediastinal, caudal mediastinal, right bronchial, left bronchial and cranial tracheobronchial) showed a scores of between 4 and 14 in animals infected with *M. bovis* AF2122/97; animals infected with *M. tuberculosis* H37Rv had a score of 1; animals infected with *M. tuberculosis* BTB1558 had scores ranging of between 0 and 2; statistically significant differences were observed only between animals infected with *M. bovis* AF2122/97 and those infected with *M. tuberculosis* BTB1558. Extra-thoracic lymph nodes from the head and neck region (right and left lateral retropharyngeal, right and left medial retropharyngeal, right and left parotid and right and left submandibular) showed scores of between 0 and 7 in animals infected with *M. bovis* AF2122/97; scores of between 0 and 2 in animals infected with *M. tuberculosis* BTB1558; animals infected with *M. tuberculosis* H37Rv did not show any gross visible lesions in extra-thoracic lymph nodes. No statistically significant differences were observed between the three groups of infected animals in the lesions observed in these nodes. No lesions were found in the tonsils in any group. All animals showed TB-like gross lesions in at least one organ.

**Figure 3:**
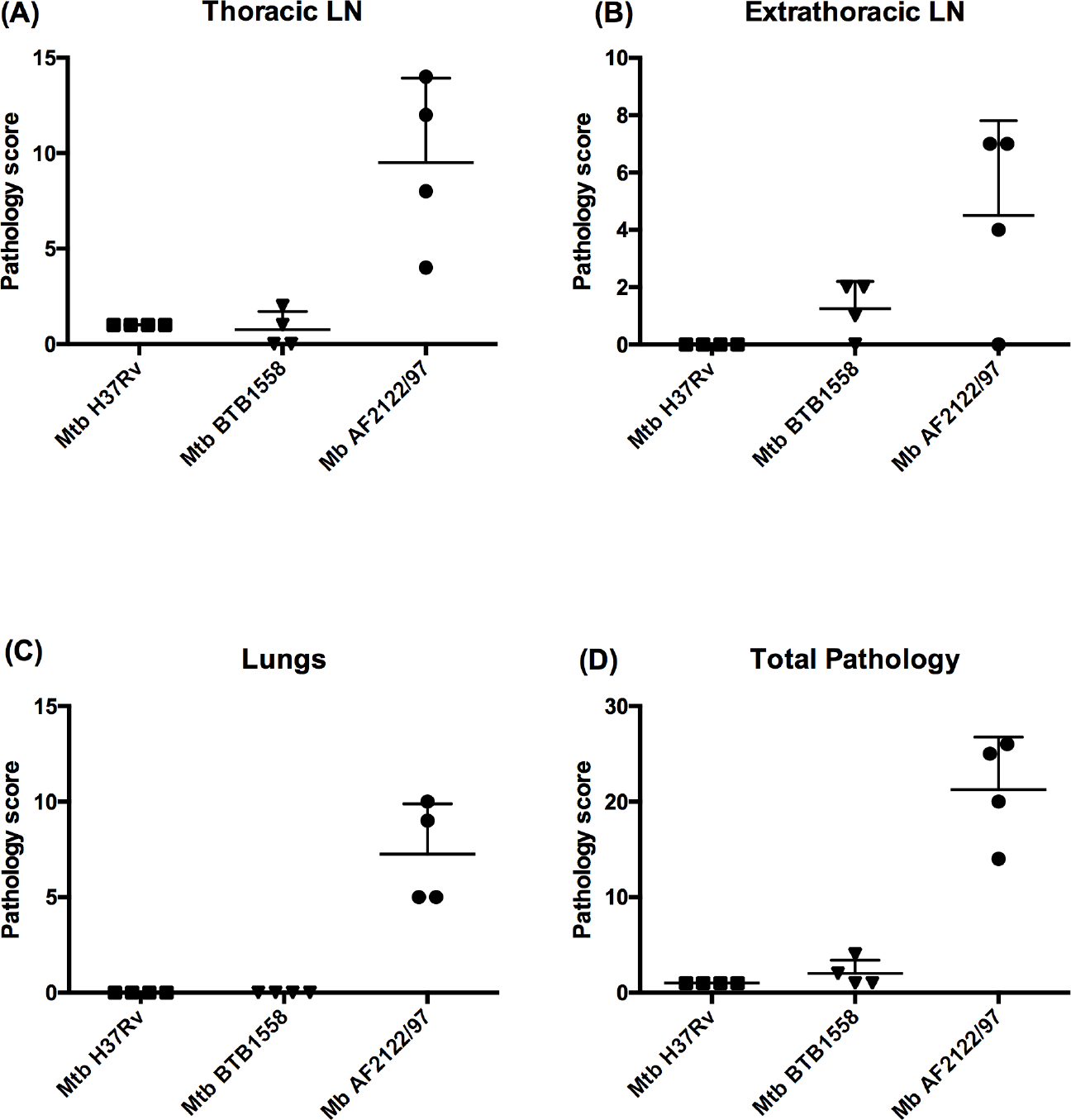
Pathology scores across *M. tuberculosis* H37Rv, *M. tuberculosis* BTB1558 or *M. bovis* AF2122/97 infected cattle. Pathology scores in the thoracic (A), extra thoracic lymph nodes (LNs) (B), and lungs (C) of animals infected with *M. tuberculosis* H37Rv (squares); *M. tuberculosis* BTB1558 (triangles), or *M. bovis* AF2122/97 (circles). Total gross pathology score is shown in (D). Data for each time point is presented as the mean response ± SEM. This difference was statistically significant in the lungs (p 0.0052) and in the total pathology score (ρ 0.0105).

**Figure 4:**
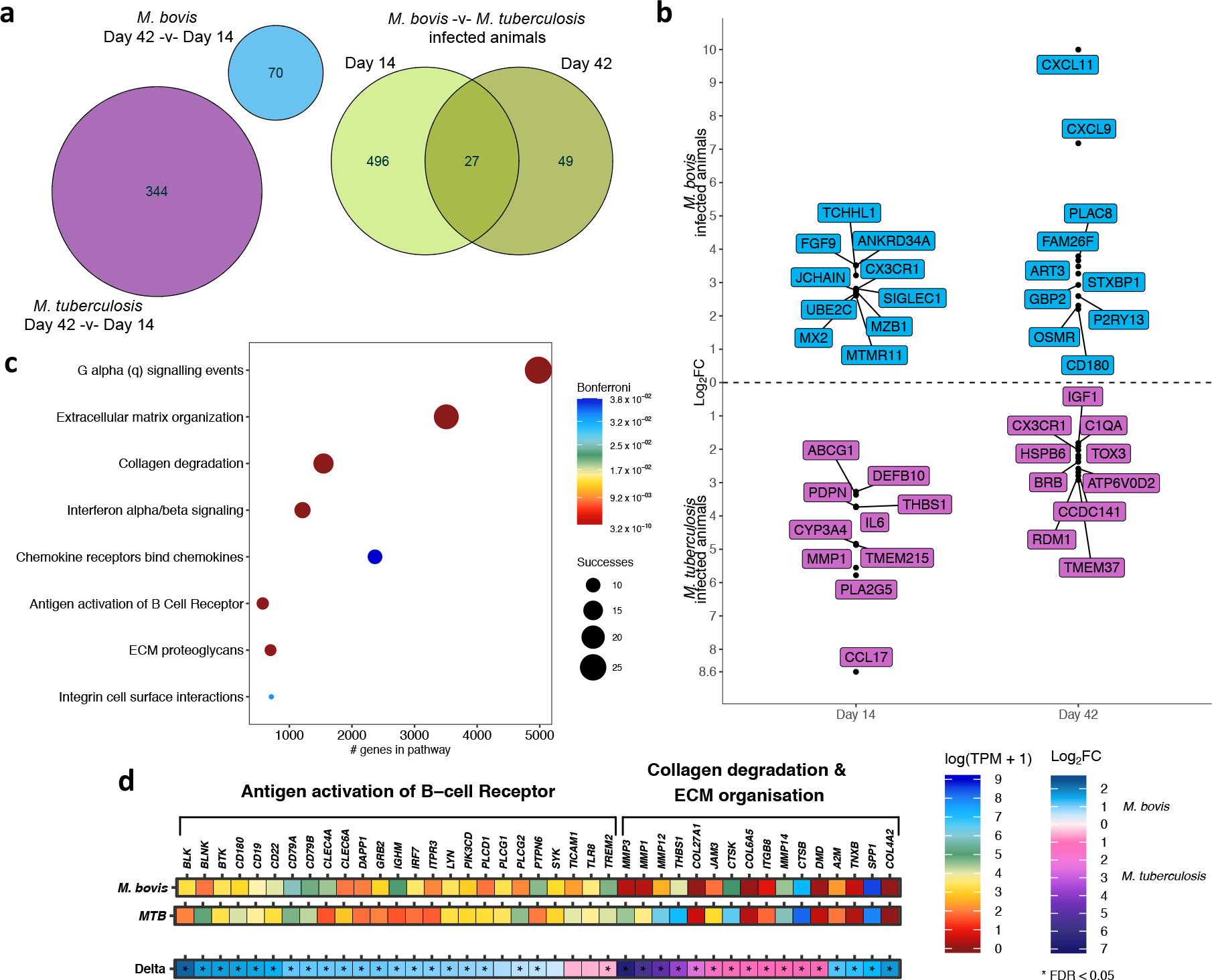
Transcriptome analysis of unstimulated whole blood after *M. bovis* AF2122/97 or *M. tuberculosis* H37Rv infection. (A) The number of differentially expressed genes in unstimulated whole blood for *M. bovis-* (blue) and *M. tuberculosis-* (purple) infected animals at day 42-v-day 14 post infection (−1 > log_2_FC < 1, FDR < 0.05). The green Venn diagram depicts the overlap of differentially expressed genes from the direct comparison of unstimulated whole blood samples from *M. bovis* and *M. tuberculosis* infected animals at day 14 and day 42 post infection (−1 > log_2_FC < 1, F DR < 0.05). (B) The top 10 differentially expressed genes between *M. bovis-* (blue) and *M. tuberculosis-* (purple) infected animals at day 14-and day 42-post infection (FDR < 0.05). The graph depicts positive relative log_2_ fold change values where a gene that shows increased expression in *M. bovis* infected animals is relative to its expression in *M. tuberculosis* infected animals and vice versa. (C) Pathway enrichment analysis results for the list of 523 differentially expressed genes between *M. bovis* and *M. tuberculosis* infected animals at day 14-post infection. The graph depicts the enrichment of each pathway in the differentially expressed gene list based on the SIGORA successes metric (circle size) and the number of genes annotated within the pathway (“#genes in pathway”) while the colour bar depicts the significance of the association (Bonferroni < 0.05). (D) The relative expression (transcripts per million, TPM, “log(TPM +1)”) and the relative change in expression (log_2_ fold change (“Log_2_FC”)) of the genes belonging to the Antigen activation of B cell Receptor (R-HSA-983695), Collagen degradation (R-HSA-1442490) and Extracellular matrix (“ECM”) organization (R-HSA-1474244) pathways enriched for the comparison of *M. bovis* and *M. tuberculosis* infected animals at day 14 post infection. Genes that pass multiple hypothesis testing are denoted with an asterisk (Benjamini-Hochberg, FDR < 0.05) (*).

### Culture of M. bovis and M. tuberculosis from processed samples

The presence of bacteria in the harvested tissue samples was investigated by semi-quantitative culture (Table 1). It was possible to culture the respective infecting strain from at least one organ from all experimental animals. However, the number of CFU/ml was usually low for animals infected with either of the two *M. tuberculosis* strains, with zero bacterial counts recorded in the lung or lung lymph nodes for all eight animals infected with either strain of *M. tuberculosis*. Respiratory lymph nodes were mostly affected, with all 12 animals having bacterial counts in this organ system (Table 1).

**Table 1:** Bacteriology of investigated organs.

| Strain | *M. bovis* AF2122/97 |  |  |  | *M. tuberculosis* H37Rv |  |  |  | *M. tuberculosis* BTB1558 |  |  |  |
| --- | --- | --- | --- | --- | --- | --- | --- | --- | --- | --- | --- | --- |
| Animal ID | 1217 | 1221 | 1222 | 1224 | 1201 | 1203 | 1213 | 1211 | 1202 | 1207 | 1209 | 1215 |
| Head LN ¶ | 0/8 * | 2/8 | 6/8 | 5/8 | 0/8 | 1/8 | 2/6 | 0/8 | 2/7 | 2/8 | 1/8 | 1/8 |
| Resp LN ¶ | 5/5 | 5/5 | 4/5 | 5/5 | 1/5 | 2/5 | 2/5 | 3/5 | 3/5 | 1/5 | 4/4 | 2/5 |
| Lung* | 2/2 | 1/2 | 1/1 | 2/2 | 0/1 | 0/1 | 0/1 | 0/1 | 0/1 | 0/1 | 0/1 | 0/1 |

¶LN = lymph nodes; see Materials and Methods for details of tissues examined.
\*Number of tissues sampled in each organ system per challenged animal that were confirmed as culture positive with the respective infecting strain; hence 0/8 in head lymph nodes means no positive cultures from 8 samples taken.

### Histopathology

In H&E stained sections, four stages of granulomas were identified as previously described ^39,50^. Briefly, Stage I (initial) granulomas comprised clusters of epithelioid macrophages, low numbers of neutrophils and occasional Langhans’ multinucleated giant cells (MNGCs). Stage II (solid) granulomas were more regular in shape and surrounded by a thin and incomplete capsule. The cellular composition was primarily epithelioid macrophages, with Langhans’ MNGCs present and some infiltration of lymphocytes and neutrophils. Necrosis was minimal or not present. Stage III (necrotic) granulomas were all fully encapsulated with central areas of necrosis. The necrotic centres were surrounded by epithelioid macrophages and Langhans’ MNGCs, and a peripheral zone of macrophages, clustered lymphocytes and isolated neutrophils extended to the fibrotic capsule. Stage IV (mineralised) granulomas were completely surrounded by a thick fibrous capsule and displayed central areas of caseous necrosis with extensive mineralization. The central necrosis was surrounded by epithelioid macrophages and Langhans’ MNGCs cells with a peripheral zone of macrophages and dense clusters of lymphocytes just inside the fibrous capsule. Granulomas were frequently multicentric, with several granulomas coalescing.

The number of granulomas of each stage in each tissue was variable (Table 2). Most of the histopathological lesions were observed in the thoracic lymph nodes with only two animals from BTB1558 and two others from the *M. bovis* AF2122/97 group showing granulomas in extrathoracic lymph nodes. The number of granulomas observed in tissues from animals infected with *M. bovis* AF2122/97 was significantly higher than the small number of granulomas observed in animals infected with either strain of *M. tuberculosis*. Moreover, the majority of granulomas observed in both H37Rv and BTB1558 infected animals were classed as stage I, with a few stage II granulomas, while *M. bovis* AF2122/97 infected animals showed granulomas of all development stages (I-IV) (Table 2). AFBs were identified in at least one ZN stained tissue section from every animal (data not shown)

**Table 2:** Granuloma presentations in infected cattle

| **Strain** | **Stage I<sup>¶</sup>** | **Stage II** | **Stage III** | **Stage IV** |
| --- | --- | --- | --- | --- |
| *M. tuberculosis* H37Rv | 31 | 2 | 0 | 0 |
| *M. tuberculosis* H37Rv | 20 | 5 | 0 | 0 |
| *M. bovis* AF2122/97 | 74 | 117 | 51 | 81 |

¶Sections were scored for granuloma stages: stage I (initial), stage II (solid), stage III (necrotic) and stage IV (mineralised). Granuloma development in both *M. tuberculosis* strains is significantly different from *M. bovis* AF2122/97, $P<0.001$ (X-sq for trend)

### *Transcriptome analysis of* M. bovis *AF2122/97 and* M. tuberculosis *H37Rv infected animals*

As the peripheral immune responses were broadly similar across all groups, yet pathological examination revealed significant differences, we used transcriptomic analysis of stimulated and non-stimulated whole blood samples as an unbiased tool to identifying global peripheral blood markers that would correlate with the pathological outcomes. We chose to analyse just the *M. bovis* AF2122/97 and *M. tuberculosis* H37Rv groups as they presented the extremes in terms of immune responses and pathological presentations. Blood samples cultured with medium alone (negative control) or with PPD-B from *M. bovis* AF2122/97 (n=4) and *M. tuberculosis* H37Rv (n=4) infection groups at days 14 and 42 were selected for analysis. Strand-specific RNA-Seq libraries (*n* =32) were prepared from these blood samples and after sequencing on an Illumina HiSeq 2500, quality-control and filtering of sequencing reads yielded a mean of 14,981,780 paired-reads per individual library (2 x 100 nucleotides); these data satisfy previously defined criteria for RNA-Seq experiments with respect to sequencing depth ^51–53^. Alignment of filtered paired-end reads to the *B. taurus* reference genome UMD3.1.1 yielded a mean of ~13.5 million read pairs (~90.5%) mapping to unique locations per library. Gene count summarisation resulted in an average of 59% of read pairs being assigned to *B. taurus* reference genome annotations based on strict sense strand and counting specifications. Gene filtering resulted in 17,663 sense genes (57.5% of all *B. taurus* reference genes in the RefSeq annotations) that were suitable for differential expression analysis. Multidimensional scaling analysis amongst the 32 sequencing libraries using the filtered and normalised gene counts (*n* = 17,663 genes) revealed that PPD-B stimulation was the largest discriminator that placed the samples into two distinct groups (Figure S1).

### Differential gene expression in unstimulated whole blood from M. bovis AF2122/97 and M. tuberculosis H37Rv infected animals

Pairwise analysis of the transcriptome from unstimulated bovine whole blood samples at day 42 versus day 14 revealed increases in the number of differentially expressed (DE) genes at day 42 post infection with either *M. bovis* AF2122/97 or *M. tuberculosis* H37Rv (Figure 5A), with a greater number of genes found differentially expressed in *M. tuberculosis* H37Rv infected animals at day 42, respectively (344 vs. 70 genes).

**Figure 5:**
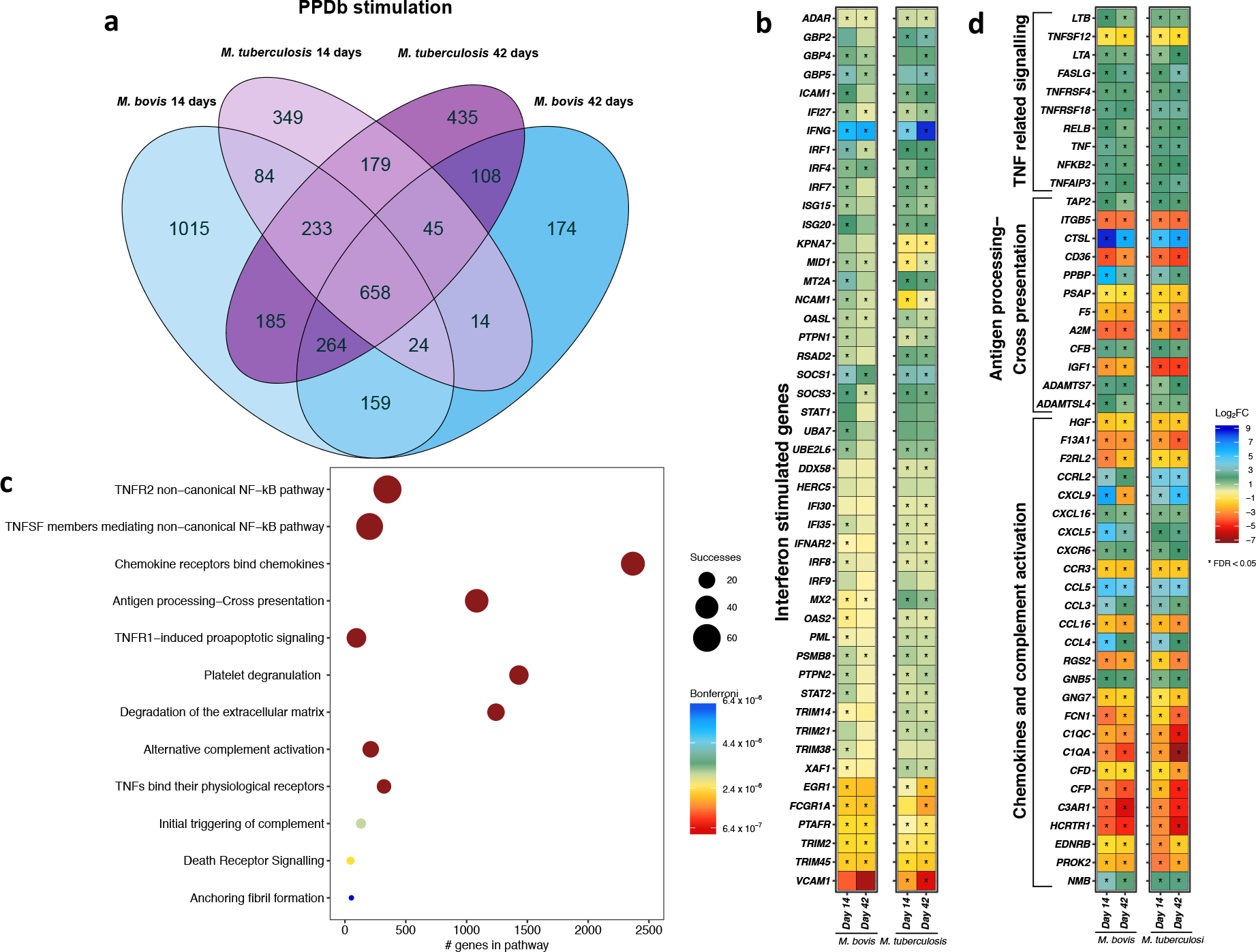
Comparative transcriptome analysis of unstimulated vs. PPD-B stimulated whole blood from *M. bovis* AF2122/97 or *M. tuberculosis* H37Rv infected cattle. (A) The overlap of differentially expressed genes from the comparison of unstimulated whole blood samples and PPD-B stimulated whole blood samples for *M. bovis* AF2122/97 infected animals at day
and day 42 (blue) and *M. tuberculosis* H37Rv infected animals at day 14 and day 42 (purple) (−1 > log2FC < 1, FDR < 0.05). (B) The relative expression of Interferon stimulated genes (R-HSA-877300) from the comparison of unstimulated whole blood samples and PPD-B stimulated whole blood samples for *M. bovis* AF2122/97 infected animals at day 14 and day 42 (blue) and *M. tuberculosis* H37Rv infected animals at day 14 and day 42 (purple). Genes that pass multiple hypothesis testing are denoted with an asterisk (Benjamini-Hochberg, FDR < 0.05) (*). (C) Pathway enrichment analysis results for 658 genes that are significantly differentially expressed (−1 > log_2_FC < 1, FDR < 0.05) in PPD-B stimulated whole blood samples from both *M. bovis* AF2122/97-and *M. tuberculosis* H37Rv-infected animals at both 14 and 42 days post infection in comparison to unstimulated whole blood samples. The graph depicts the association of each pathway with the differentially expressed gene list based on the SIGORA successes metric (circle size) and the number of genes annotated within the pathway (“#genes in pathway”) while the colour bar depicts the significance of the association (Bonferroni < 0.05). (D) The relative expression (transcripts per million, TPM, “log(TPM +1)”) and the relative change in expression (log_2_ fold change (“Log_2_FC”)) of 658 genes that are significantly differentially expressed (-1 > log_2_FC < 1, FDR < 0.05) in PPD-B stimulated whole blood samples from both *M. bovis* AF2122/97-and *M. tuberculosis* H37Rv-infected animals at both 14 and 42 days post infection in comparison to unstimulated whole blood samples. The genes are associated with pathways related to TNF signalling (R-HSA-5668541, R-HSA-5676594, R-HSA-5357786, R-HSA-5669034), Antigen processing-Cross presentation (R-HSA-1236975) and Chemokines and complement activation (R-HSA-380108, R-HSA-173736). Genes that pass multiple hypothesis testing are denoted with an asterisk (Benjamini-Hochberg, FDR < 0.05) (*).

Direct comparison of unstimulated whole blood samples between *M. bovis* AF2122/97 and *M. tuberculosis* H37Rv infected animals at day 14 and again at day 42 revealed 523 and 76 DE gene respectively, with 27 of these identified as DE at both time-points (Figure 5A, green) between the two infection models. Amongst the top 10 DE genes upregulated in *M. bovis* AF2122/97 infected animals (or conversely, downregulated in *M. tuberculosis* H37Rv animals) included those encoding: the macrophage restricted cell surface receptor *SIGLEC1* that is known to be involved in pathogen uptake, antigen presentation and lymphocyte proliferation ^54^; the CD4-coreceptor and fractalkine receptor *CX3CR1*, which has been linked to tuberculosis susceptibility and the impairment of macrophage and dendritic cell migration ^55,56^; and Interferon-Regulated Resistance GTP-Binding Protein *(MX1)* at day 14 (Figure 5B). At day 42 the expression of the following genes was at relatively higher level in *M. bovis* AF2122/97 infected animals than in *M. tuberculosis* H37Rv infected animals (Figure 5B): *CXCL9*, a previously described potential TB biomarker ^50,57^; mediator of mycobacterial-induced cytokine production in macrophages *CD180* ^58^; and the chemokine *CXCL11*.

T-cell chemotactic factor *CCL17* had the highest log-fold change in *M. tuberculosis* H37Rv infected samples in comparison to *M. bovis* AF2122/97 infections at day 14. Genes also expressed to a higher level in *M. tuberculosis* infected animals at day 14 were: the defensin *DEFB10;* platelet aggregation inducing factor *PDPN*, which is expressed on macrophages and epithelioid cells within the tuberculous granuloma ^59^; macrophage lipid export complex member *ABCG1;* and *IL6* (in contrast with the ELISA data from the same time point, Figure 2). At day 42 post infection the fractalkine receptor *CX3CR1*, a marker of Th1 stage differentiation during tuberculosis ^60^, and the V-ATPase subunit gene *ATP6V0D2* were higher in *M. tuberculosis* H37Rv infected samples (Figure 5B).

Pathway enrichment analysis revealed pathways such as *Extracellular matrix organization* (R-HSA-1474244), *Collagen degradation* (R-HSA-1442490), *Interferon alpha/beta signaling* (R-HSA-909733) and B cell receptor second messenger signaling (R-HSA-983695) being significantly associated with the 523 DE genes between *M. tuberculosis* and *M. bovis* AF2122/97 infected animals at day 14 (Bonferroni < 0.005) (Figure 6C, 6D). Further investigation revealed higher expression of 25/44 genes belonging to the antigen activation of B cell receptor signaling pathway (R-HSA-983695) in *M. bovis* AF2122/97 infected animals at day 14, such as membrane associated *CD19, CD80, TREM2* and second messengers *LYN*, *SYK*, *BTK*, *BLNK*, *PLCG2* along with B-cell receptor encoding genes *CD79a* and *CD79b* (Figure 6D). The enrichment of collagen degradation and extracellular matrix organization pathways in the 523 DE gene list highlighted genes of the matrix metallopeptidase (MMP) family such as *MMP1, MMP3, MMP12* and *MMP14* (Figure 6C). MMP proteins have been linked to leukocyte migration and the progression of granuloma formation during tuberculosis; *MMP1* and *MMP14* gene products are key for the destruction of collagen and alveolar destruction with an increased expression of *MMP14* gene found in the sputum of tuberculosis patients ^61,62^. Gene expression of the matrix-associated cytokine SPP1 (i.e. Osteopontin/OPN) was upregulated in *M. bovis* infected animals at day 14. SPP1 enhances IFN-γ and IL-12 production, and increased levels of SPP1 have been reported in the blood of TB patients versus controls ^63^.

Pathway enrichment analysis with the 76 common DE genes at day 42-post infection revealed no significantly associated pathways.

### Differential gene expression in PPD-B-stimulated whole blood from M. bovis AF2122/97 and M. tuberculosis H37Rv infected animals

To assess antigen-stimulated alteration in the whole blood transcriptome, blood samples were stimulated with PPD-B overnight and subsequently compared to unstimulated blood samples. PPD-B stimulation resulted in the differential expression of 2,622 and 1,586 genes at day 14 and 1,446 and 2,107 at day 42 in *M. bovis* AF2122/97 and *M. tuberculosis* H37Rv infected animals respectively in comparison to control samples (Figure 6A).

Firstly, a “core” response to PPD-B stimulation amounting to 658 genes was identified, representing genes that were consistently DE regardless of infectious agent or time post infection (Figure 6A). As expected, *IFNY* was strongly upregulated in stimulated blood samples at both time points in both *M. bovis* AF2122/97 and *M. tuberculosis* H37Rv infected animals (Figure 6B). Furthermore, 47/152 genes from the *Interferon gamma signaling* pathway (R-HSA-877300) were significantly differentially expressed (−0.75 < log_2_FC > 0.75) across the comparative groups and time points (Figure 6B). Pathway enrichment analysis of the 658 core response genes revealed significantly associated pathways such as those involved in TNF-signaling (R-HSA-5668541, R-HSA-5676594, R-HSA-5357786, R-HSA-5669034), *Chemokine receptors bind chemokines* (R-HSA-380108), *Antigen processing-Cross presentation* (R-HSA-1236975) and *Initial triggering of complement* (R-HSA-173736) (Figure 6C). The change in expression of the genes within these pathways can be seen in Figure 6D; an overall downwards trend in genes encoding complement related factors was found (*e.g. C1Q1, C1QA, CFD* and *CFP*) along with strong upregulation of TNF-signaling related factors *(e.g. TNF, NFKB2, TNFAIP3* and lymphotoxins alpha and beta *LTA/LTB)* upon PPD-B stimulation of whole blood samples (Figure 6D).

Second, there are 159 and 179 DE genes in either *M. bovis* AF2122/97 or *M. tuberculosis* H37Rv infected animals at both time points (Figure 6A). These 338 genes represent a divergence in response to PPD-B stimulation between the two infection groups and the top ranking genes amongst them are presented in Figure S2 (log_2_FC ratio). The increased expression of *CCL17, DEFB10* and matrix metalloproteinase *MMP12* with the decreased expression of *bactericidal permeability increasing protein BPI, CD164* and IL5 receptor *IL5RA* can differentiate *M. bovis* AF2122/97 infected animals from *M. tuberculosis* H37Rv infected animals at both day 14 and day 42 post infection upon PPD-B stimulation. Conversely, the increased expression of interferon inducible dyamin *MX2* at day 14 and cytokine receptors *IL22RA2* and *XCR1* at day 42 post infection, and decreased expression of T-cell regulator *TNFSF18* at day 14-and *TLR5*, defensin *DEFB5* and V-ATPase subunit gene *ATP6V0D2* at day 42-post infection, distinguished *M. tuberculosis* H37Rv infected from *M. bovis* AF2122/97 infected animals upon PPD-B stimulation.

### miR-155 analysis

miR-155 has been identified as a potential biomarker of disease development and/or severity in cattle infected with *M. bovis* ^49^. To explore its utility, we analysed the expression of miR-155 in whole blood from *M. bovis* AF2122/97 and *M. tuberculosis* H37Rv infected cattle in PPD-B stimulated vs. unstimulated whole blood over the infection time course using RT-qPCR (Figure S3). The animals infected with *M. tuberculosis* H37Rv showed a steady increase in expression of miR-155 after PPD-B stimulation over the course of infection, while *M. bovis* infected animals showing a greater baseline expression prior to infection and hence a more modest increase over the time course. While a single miRNA may lack specificity as a biomarker, these results support the potential of miR-155 as a part of a biomarker panel to assess infection with tubercle bacilli.

## Discussion

This work set out to explore whether a recent isolate of *M. tuberculosis*, recovered from a TB lesion identified in an Ethiopian bull, would trigger a similar immunological and pathological response to the *M. tuberculosis* H37Rv reference strain when used to experimentally infect cattle. This work sought to build on our previous study that had shown *M. tuberculosis* H37Rv to be attenuated in the bovine host but using a recent clinical isolate to address issues around possible laboratory-adaptation of the H37Rv strain that may have reduced its virulence in the bovine model. Furthermore we wished to see whether the application of transcriptomics would reveal potential biomarkers to differentiate between the initial stages of a progressive, active *M. bovis* infection and a more quiescent, latent infection with *M. tuberculosis*.

Our results showed that both *M. tuberculosis* BTB1558 and *M. tuberculosis* H37Rv were considerably attenuated in the bovine host when compared to *M. bovis* AF2122/97. This is despite the fact that animals infected with either *M. tuberculosis* strain were left to progress for 10 weeks, while the *M. bovis* infected animals were culled 6 weeks after infection. *M. bovis* induced a greater level of pathology than either of the *M. tuberculosis* strains. Although *M. tuberculosis* BTB1558 appeared to induce a slightly greater level of pathology in head lymph nodes than *M. tuberculosis* H37Rv, this difference was not significant; no difference was detected in the level of pathology induced by either of the *M. tuberculosis* strains in respiratory lymph nodes or lungs. The apparent arrest of *M. tuberculosis* granulomas in stages I and II, compared to infection with *M. bovis* that produced lesions from stages I-IV is another striking difference in infection outcome. The bacteriological culture results showed that both strains of *M. tuberculosis* persisted in cattle, at least over the 10 weeks of the infection time course.

The kinetics and magnitude of peripheral blood responses across all three infection groups were broadly similar over the initial 6 week phase of infection. As analysis of antigen-induced peripheral blood cytokine responses failed to reflect the distinct pathological presentations between the *M. bovis* AF2122/97 and *M. tuberculosis* groups, we applied RNA-Seq transcriptomics of whole blood in an attempt to identify biomarkers that would better distinguish the groups, focusing on the *M. bovis* AF2122/97 and *M. tuberculosis* H37Rv groups. This analysis revealed discrete genes and pathways that distinguished the groups and were indicative of accelerated disease development in the *M. bovis* AF2122/97 infected animals, and included previously described biomarkers of disease progression or susceptibility such as MMPs, CX3CR1, CXCL9 and SSP1/OPN ^50,57,60,62,63^. These changes in peripheral gene expression over the course on infection provide biomarker candidates of disease progression for validation in new studies.

Classic experiments from the late 19^th^ century by Smith, Koch and von Behring showed that bovine and human tubercle bacilli showed distinct virulence in animal models, and in particular that the human bacillus was attenuated when used to infect cattle. Our work has recapitulated these findings, using both the standard *M. tuberculosis* H37Rv strain as well as an *M. tuberculosis* isolate (BTB1558) recovered from a bovine lesion. The attenuation shown by the *M. tuberculosis* BTB1558 strain in this experimental model, compared to that seen in the field situation in Ethiopia, therefore appears likely due to increased susceptibility of the affected animal. It should be noted however that from a human and animal health perspective, it is possible that cattle infected with *M. tuberculosis* may still be transmissible and shed bacilli (e.g. through nasal secretions, aerosol from the lungs or through milk) albeit at low levels. Such scenarios are more probable in countries where the case rates of active human TB are high, interaction between humans and cattle are frequent, and consumption of raw milk common.

While the outcome of infection with *M. tuberculosis* in humans is likely more a spectrum than a bipolar, active vs. latent, state ^64,65^, infection with *M. tuberculosis* in cattle could be indicative of a latent infection being established, as compared to an active disease status upon *M. bovis* infection. The bovine infection model may therefore offer a tractable experimental system in which to explore the reactivation of *M. tuberculosis* infection and to define prognostic biomarkers of the development of active disease. Furthermore, infection of cattle with *M. tuberculosis* to establish latent infection may provide a tractable outbred model in which to explore post-exposure vaccination strategies.

In conclusion, our work has shown that *M. tuberculosis* isolates, whether the H37Rv type strain or a recent isolate, are attenuated for virulence in a bovine infection model as compared to *M. bovis*. This work provides further evidence of the distinct host preference of tubercle bacilli as a basis to explore the molecular basis of virulence in *M. bovis* as compared to *M. tuberculosis*, and also offers a model in which to explore the reactivation of latent TB infection.

## Acknowledgements

The authors are grateful to the members of the scientific and animal staff of the Plate-Forme d’Infectiologie Expérimental, PFIE, INRA, 37380, Nouzilly, France, especially to the study managers Céline Barc and Philippe Bernardet and animal technicians Olivier Boulesteix and Joël Moreau.

---

**Figure S1:**
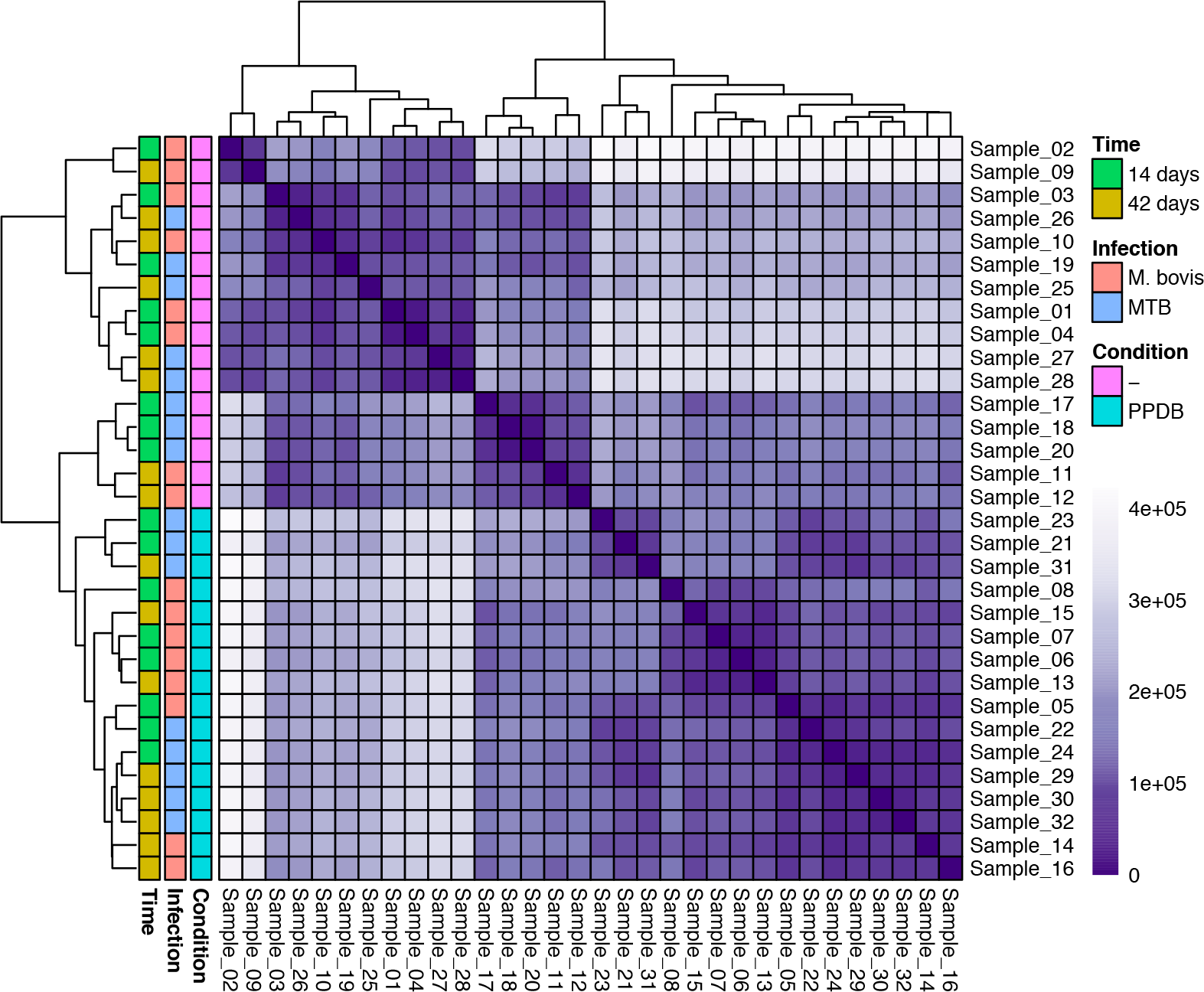
Genome-wide gene expression correlation. Genome-wide gene expression correlation between the 32 study samples pertaining to whole blood samples (unstimulated (“-“) or PPDb-stimulated (“PPDB”)) from cattle infected with either M. bovis AF2122/97 (“M. bovis”) or *M. tuberculosis* H37Rv (“MTB”) 14 days or 42 days (“Time”) post infection. Samples are clustered using Euclidean distance and coloured bars on the left of the plot denote the variables time point (“Time”), infection status (“Infection”) and stimulation status (“Condition”) for each of the 32 samples.

**Figure S2:**
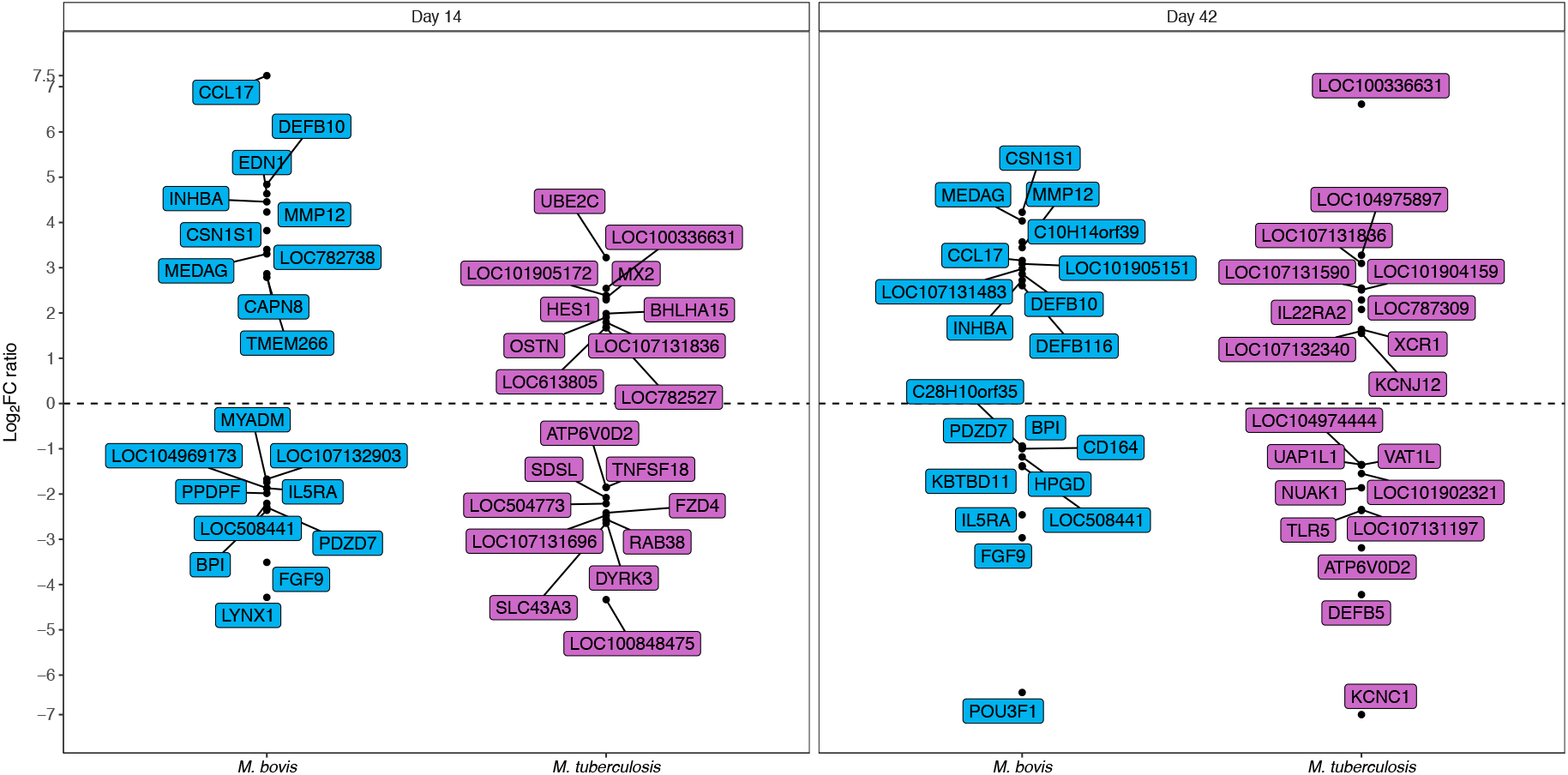
Top DE genes in unstimulated vs. stimulated whole blood between *M. bovis* AF2122/97 and *M. tuberculosis* H37Rv infected animals. The top 10 upregulated and top 10 downregulated differentially expressed genes in whole blood samples derived from *M. bovis* AF2122/97 versus *M. tuberculosis* H37Rv infected animals and stimulated with PPD-B at day 14 and day 42 post infection. The change in gene expression from the comparison of stimulated blood to unstimulated blood at each time point for either *M. bovis* AF2122/97 or *M. tuberculosis* H37Rv infected animals was used to calculate log_2_FC ratio (i.e. expression of gene X in *M. bovis* AF2122/97 infected animals at day 14-post infection divided by expression of gene X in *M. tuberculosis* H37Rv infected animals at day 14-post infection).

**Figure S3:**
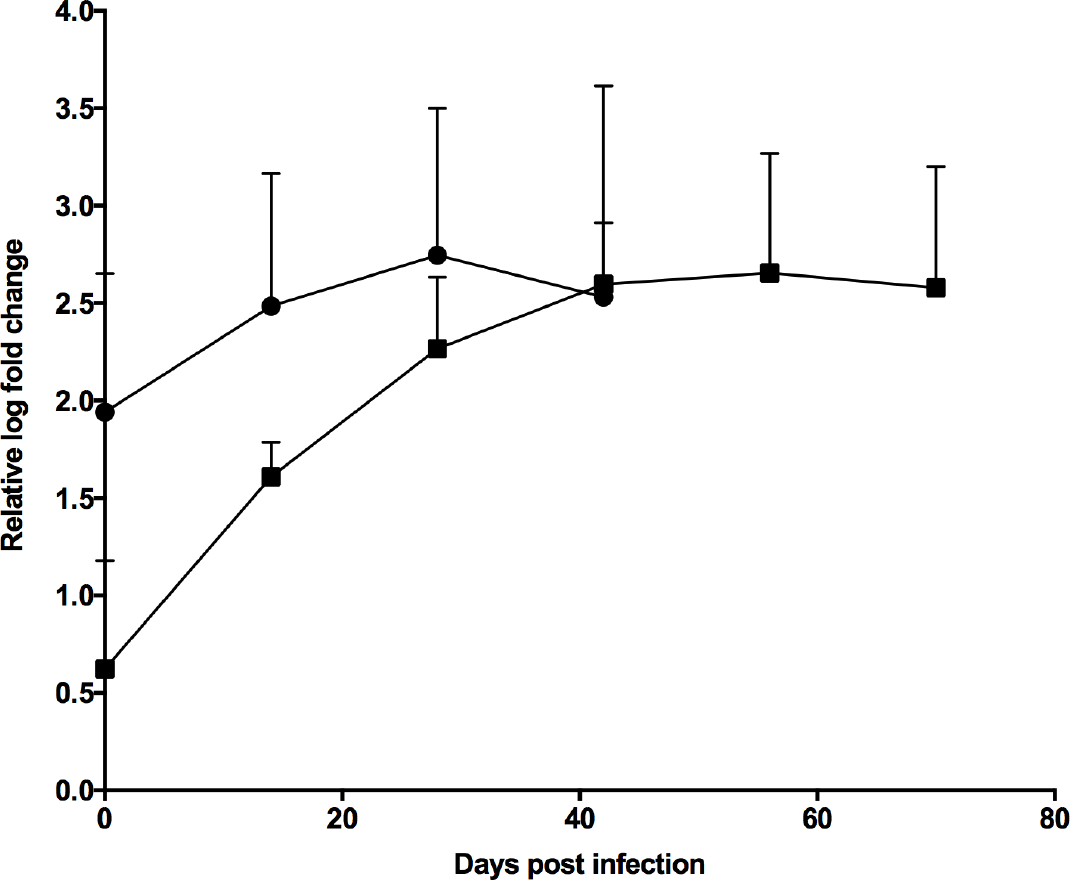
miR-155 analysis across *M. bovis* AF2122/97 and *M. tuberculosis* H37Rv infected animals. The level of miR-155 in PPD-B stimulated vs. unstimulated whole blood from *M. bovis* AF2122/97 (circles) and *M. tuberculosis* H37Rv (squares) infected cattle was assessed over the infection time course using RT-qPCR with Exiqon miRCURY UniRT miRNA hsa-miR-155-5p primers.

